# Transdiagnostic characterization of neuropsychiatric disorders by hyperexcitation-induced immaturity

**DOI:** 10.1101/376962

**Authors:** Tomoyuki Murano, Hideo Hagihara, Katsunori Tajinda, Mitsuyuki Matsumoto, Tsuyoshi Miyakawa

## Abstract

Biomarkers are needed to improve the diagnosis of neuropsychiatric disorders. Promising candidates are imbalance of excitation and inhibition in the brain, and maturation abnormalities. Here, we characterized different disease conditions by mapping changes in the expression patterns of maturation-related genes whose expression was altered by experimental neural hyperexcitation in published studies. This revealed two gene expression patterns: decreases in maturity markers and increases in immaturity markers. These two groups of genes were characterized by the overrepresentation of genes related to synaptic function and chromosomal modification, respectively. We used these two groups in a transdiagnostic analysis of 80 disease datasets for eight neuropsychiatric disorders and 12 datasets from corresponding animal models, and found that transcriptomic pseudoimmaturity inducible by neural hyperexcitation is shared by multiple neuropsychiatric disorders, such as schizophrenia, Alzheimer disorders, and ALS. Our results indicate that this endophenotype serve as a basis for transdiagnostic characterization of these disorders.

## Introduction

Neuropsychiatric disorders—such as schizophrenia, bipolar disorder, major depression disorders, and autism spectrum disorder—are common, with over a third of the population in most countries being diagnosed with at least one such disorder at some point in their life^1^. Almost all neuropsychiatric disorders are currently classified mainly on the basis of clinical signs and symptoms. However, there is evidence that patients with different clinical diagnoses share similar biological features, such as genetic mutations, molecular expression, or brain activity^2–6^. Recently, psychiatry has undergone a tectonic shift to incorporate the concepts of modern biology. There have been recent attempts to reclassify psychiatric disorders according to biological domains (e.g. genes, neural circuits, behavior), such as through the Research Domain Criteria (RDoC) initiative^7^. Therefore, identifying appropriate biomarkers which can be used for transdiagnostic assessment of neuropsychiatric disorders is essential for better classification of these diseases and understanding of their biological basis.

Using coexpression network analysis, a recent study revealed that cross-disorder gene expression overlaps could be used to characterize five major neuropsychiatric disorders^8^. Some of these overlapping gene groups were biologically well-characterized by Gene Ontology enrichment or cell-type specificity, but the biological properties of other gene groups were rather unclear. Thus, non-biased coexpression network analyses do not necessarily detect modules that extract the biological features of neuropsychiatric disorders. Then, to achieve better characterization of neuropsychiatric disorders, it might be helpful to detect modules of coexpressed genes and conduct gene expression analysis based on the findings derived from researches on animal models of neuropsychiatric disorders.

To date, we have screened more than 180 strains of genetically engineered mice using a large-scale and comprehensive battery of behavioral tests, and identified several strains with abnormal behaviors related to neuropsychiatric disorders such as schizophrenia, bipolar disorder, and intellectual disability^9^. We discovered common endophenotypes in the brains of multiple strains of these genetically engineered mice with behavioral abnormalities. We termed one such endophenotype in the hippocampus of adult mice the “immature dentate gyrus (iDG)” phenotype^10–13^. In this phenotype, the molecular and electrophysiological properties of adult DG neurons in the genetically engineered mice were similar to those of immature DG neurons in typically developing infants. For example, the expression of calbindin, a marker of maturity in DG neurons, was decreased and the expression of calretinin, a marker of immaturity, was increased^10–15^. Similar molecular changes to some of those found in mice with iDG have been observed in the postmortem brains of patients with schizophrenia^16^, bipolar disorder^16^, and epilepsy^17–19^. Furthermore, there is growing evidence that changes in molecular markers of pseudoimmaturity are also observed in other brain areas of patients with schizophrenia^20–28^, bipolar disorder^26^, autism^26^, and alcoholism^29^. Therefore, we proposed that pseudoimmaturity of the brain could potentially be a useful transdiagnostic biomarker^9^.

Pseudoimmaturity of the brain can be induced in adulthood. Previously, we found that chronic fluoxetine treatment reverses the maturation status of DG neurons in adult wild-type mice, a phenomenon that we termed “dematuration”^30, 31^. Likewise, recent studies suggest that several maturation-related genes and electrophysiological properties in the DG of wild-type adult mice assume an immature-like status after treatment with pilocarpine or electroconvulsive stimulation^16, 32^. As mentioned above, an iDG-like phenotype has been found in patients with epilepsy^17–19^. Therefore, we hypothesized that the hyperexcitation of neurons may be a cause of pseudoimmaturity of the brain in adulthood.

Some studies suggest that hyperexcitation of neurons may underlie abnormalities related to certain types of neuropsychiatric disorders. Individuals with epilepsy are at increased risk of developing schizophrenia, and vice versa^33, 34^, and patients with epilepsy can also display psychotic symptoms that resemble those found in patients with schizophrenia^35^. Imbalances in excitatory and inhibitory brain circuitry have been proposed to be involved in the pathogenesis and pathophysiology of schizophrenia^36–39^. Hyperactive action-potential firing has also been observed in hippocampal granule-cell-like neurons derived from induced pluripotent stem cells (iPSCs) of patients with bipolar disorder^40^. Recent studies suggested that human patients with Alzheimer’s disease and temporal lobe epilepsy may harbor common underlying mechanisms^17,41–43^. Considering these findings, we hypothesized that the immature-like gene expression patterns induced by neural hyperexcitation may overlap with the abnormal gene expression patterns in the brains of patients with neuropsychiatric disorders and the related animal models. If this is the case, we hypothesized that this overlap can be used to perform transdiagnostic characterization of neuropsychiatric disorders.

To test this hypothesis, we first performed a meta-analysis of microarray datasets, comparing the changes in gene expression in rat DG after seizure induction with the differences in gene expression in infant mice versus adult mice. To assess consistency across species, we also conducted a similar comparison with human fetal hippocampus. The overlap between gene sets was estimated using the Running Fisher test^44^, which is a nonparametric rank-based statistical method developed by BSCE. This method enables us to statistically assess the pairwise correlations between any two datasets, including datasets from different species and organs^28, 29, 45, 46^. The gene expression patterns in rat DG after seizure induction significantly overlapped with those specific to immature mouse DG and also with those specific to early-stage human fetal hippocampus. From the set of overlapping genes, we defined two groups; maturity-marker genes and immaturity-marker genes that are inducible by neural hyperexcitation. We assessed the expression patterns of these two groups of maturation-related genes in 80 public gene-expression datasets derived from the postmortem brains of patients with various neuropsychiatric disorders and from neural cells derived from patient iPSCs, and in a further 12 datasets from the brains of related animal models. Through this analysis, we characterized the expression patterns of maturation-related genes that are inducible by neural hyperexcitation across different disease conditions.

## Results

### Neural hyperexcitation induces immature-like gene expression patterns in DG

To examine the developmental changes in gene expression patterns in the rodent DG, we created a microarray dataset from postnatal day 8, 11, 14, 17, 21, 25, and 29 infant mice (GSE113727) and compared it with a dataset from 33-week-old adult mice (GSE42778)^12^. Within the entire mice DG dataset, the largest overlap for changes in gene expression after pilocarpine injection was for the comparison between day 8 infant and 33-week-old adult mice (Figure S1a). We included the dataset from postnatal day 8 infant mice for subsequent analysis. The expression levels of 6552 genes were increased in the DG of infant mice compared with adult mice, whereas the expression levels of 8637 genes were decreased (absolute fold change > 1.2 and t-test *P* < 0.05). Next, we assessed the changes in gene expression induced by neural hyperexcitation in a rodent model. We obtained publicly available microarray datasets from the DG of adult rats after seizures induced by injection of pilocarpine (GSE47752)^47^. The expression levels of 7073 genes were significantly changed in the DG of epileptic-seizure rats 1 day after pilocarpine injection compared with rats treated with saline (absolute fold change > 1.2, *P* < 0.05).

To investigate whether the neuronal hyperexcitation datasets contain immature-like gene expression patterns, we assessed the overlap between the set of genes with altered expression in immature mice and the set of genes with altered expression in adult seizure-model rats, using the Running Fisher algorithm on the BaseSpace platform to determine the significance of the overlap (see Supplementary Methods for Details). We found a striking degree of similarity: 2807 genes showed changes in expression in both datasets (overlap *P* = 3.8 × 10^-11^) (Figure 1a). Among these 2807 genes, we named the 726 genes whose expression levels decreased in both datasets “hyperexcitation-induced maturity-related genes” (hiM genes (mouse): green bar in Figure 1a) and the 938 genes whose expression levels increased in both datasets “hyperexcitation-induced immaturity-related genes” (hiI genes (mouse): red bar in Figure 1a). The comprehensive gene lists of hiM and hiI genes are in Table S1 (Table S1: hiM/hiI_GeneList). The overlap for genes with positively correlated expression (red and green bars) was larger than the overlap for genes with negatively correlated expression (light and dark yellow bars), indicating that the direction of expressional changes in the two datasets are more alike than they are different. These results suggest that neuronal hyperexcitation induces a pattern of immature-like gene expression in the adult DG.

**Figure 1.**
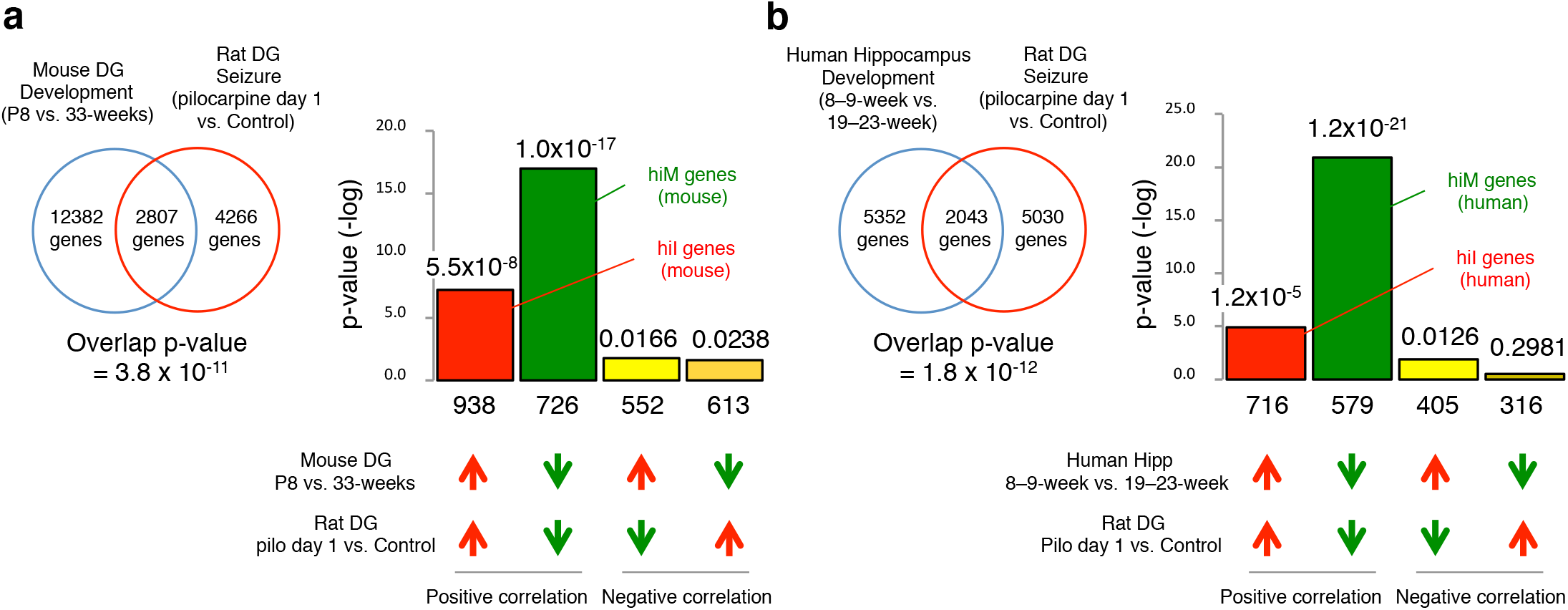
The patterns of changes in gene expression in rat DG 1 day after pilocarpine treatment compared with developmental changes in mouse DG and human hippocampus. Venn diagrams illustrating the overlap in genome-wide gene-expression changes between rat DG after seizure induction (GSE47752) and the DG of typically developing mouse infants (GSE113727; P8 infants compared with 33-week adults) (a) or the hippocampus of typically developing human fetuses (GSE25219: 19–23 week fetuses compared with 8–9 week fetuses) (b). Bar graphs illustrate the -log of the overlap *P*-values for genes upregulated (red arrows) or downregulated (green arrows) by each condition. The Bonferroni correction was used to adjust the significance level according to the number of dataset pairs (see the Methods section and Supplementary Method). Genes that were downregulated in both conditions were defined as mouse hiM genes (green bar in (a)), and genes that were upregulated in both conditions are defined as mouse hiI genes (red bar in (a)). Similarly, human hiM genes and human hiI genes were defined as the groups of genes with positive correlation between the two conditions, development and seizure (b).

Next, we compared the changes in expression during development in the human fetal hippocampus with those in rats after seizure induction to assess consistency across species. We obtained publicly available microarray datasets for the human fetal hippocampus during development (GSE25219)^48^. Within the entire fetal hippocampal dataset, the largest overlap for changes in gene expression after pilocarpine injection was for the comparison between 8–9-week fetuses and 19–23-week fetuses (Figure S1b). We again found a striking degree of similarity: 2043 genes showed changes in expression in both datasets (overlap *P* = 1.8 × 10^-12^) (Figure 1b). Among these 2043 genes, we termed the 579 genes whose expression decreased in both datasets “hiM genes (human)” (green bar in Figure 1b) and the 716 genes whose expression increased in both datasets “hiI genes (human)” (red bar in Figure 1b). The overlap for genes with positively correlated expression (red and green bars) were larger than those with negatively correlated expression (light and dark yellow bars), suggesting that, similar to the results in mice, the gene expression changes in rat DG after seizure induction are comparable to the reverse of the changes that occur as the human hippocampus develops.

### Hyperexcitation-induced maturity- and immaturity-related genes exhibit different biological properties

To characterize the biological features associated with the hiM and hiI gene groups in mouse and human, we conducted pathway enrichment analyses in BaseSpace. The 20 biogroups with the most significant overlap with hiM and hiI genes are listed in Table 1a and 1b. Among mouse hiM genes, 4 out of the top 20 biogroups are associated with synapse and channel activity (e.g., transmission of nerve impulse, synapse, and synaptic transmission) (Table 1a), whereas among human hiM genes, 6 out of the top 20 biogroups are also associated with synapse and channel activity (e.g., transmission of nerve impulse, synaptic transmission, axon, and synapse) (Table 1b).

**Table 1.** Summary of results from the pathway analyses of hiM/hiI genes. (a) The 20 biogroups with the most significant similarities to mouse hiM genes and mouse hiI genes. Green columns indicate biogroups that are related to the plasma membrane. Red columns indicate biogroups that are related to reactions in the nucleus. (b) The 20 biogroups with the most significant similarities to human hiM genes and human hiI genes.

| hiM genes (mouse) |  |  |  |
| --- | --- | --- | --- |
| Biogroup name | direction | Common genes | p-value |
| multicellular organismal signaling | down | 56 | 4.00E-23 |
| transmission of nerve impulse | down | 51 | 3.10E-19 |
| axon | down | 45 | 7.30E-19 |
| neuron differentiation | down | 58 | 4.70E-17 |
| synapse | down | 57 | 5.70E-17 |
| neuron development | down | 50 | 1.50E-16 |
| neuronal cell body | down | 45 | 5.20E-16 |
| cell junction | down | 56 | 3.00E-15 |
| cell part morphogenesis | down | 43 | 4.70E-15 |
| neuron projection development | down | 42 | 6.20E-15 |
| cell projection part | down | 56 | 6.50E-15 |
| cell morphogenesis involved in neuron differentiation | down | 35 | 2.90E-14 |
| cell morphogenesis involved in differentiation | down | 40 | 6.70E-14 |
| cellular chemical homeostasis | down | 53 | 2.50E-13 |
| cell-cell signaling | down | 44 | 2.60E-13 |
| synaptic transmission | down | 37 | 3.60E-13 |
| regulation of neurological system process | down | 31 | 1.10E-12 |
| regulation of transmission of nerve impulse | down | 30 | 1.40E-12 |
| Genes involved in Neuronal System | down | 31 | 1.70E-12 |
| cellular ion homeostasis | down | 49 | 1.80E-12 |

b
| hiM genes (human) |  |  |  |
| --- | --- | --- | --- |
| Biogroup name | direction | Common genes | p-value |
| multicellular organismal signaling | up | 87 | 4.40E-58 |
| transmission of nerve impulse | up | 83 | 1.40E-54 |
| synaptic transmission | up | 75 | 6.70E-50 |
| neuron projection | up | 70 | 2.60E-44 |
| axon | up | 42 | 2.90E-33 |
| synapse | up | 51 | 2.60E-32 |
| neuron development | up | 58 | 1.60E-30 |
| Genes involved in Neuronal System | up | 41 | 3.40E-30 |
| Genes involved in Transmission across Chemical Synapses | up | 32 | 1.50E-29 |
| regulation of neurological system process | up | 35 | 6.50E-29 |
| regulation of transmission of nerve impulse | up | 34 | 9.20E-29 |
| cell projection part | up | 53 | 1.80E-27 |
| passive transmembrane transporter activity | up | 44 | 1.80E-26 |
| ion channel activity | up | 43 | 2.20E-26 |
| cell part morphogenesis | up | 49 | 1.80E-24 |
| behavior | up | 38 | 2.90E-24 |
| single-organism behavior | up | 35 | 3.70E-24 |
| cell morphogenesis involved in neuron differentiation | up | 44 | 4.30E-24 |
| neuron projection development | up | 47 | 4.40E-24 |
| dendrite | up | 37 | 2.00E-23 |

| hiL genes (mouse) |  |  |  |
| --- | --- | --- | --- |
| Biogroup name | direction | Common genes | p-value |
| response to wounding | up | 60 | 1.10E-16 |
| positive regulation of developmental process | up | 66 | 9.30E-16 |
| cardiovascular system development | up | 61 | 7.70E-15 |
| circulatory system development | up | 61 | 7.70E-15 |
| proteinaceous extracellular matrix | up | 32 | 7.80E-15 |
| Genes involved in Cell Cycle | up | 47 | 1.00E-14 |
| extracellular matrix | up | 37 | 1.30E-14 |
| Genes involved in Cell Cycle, Mitotic | up | 42 | 6.30E-14 |
| protein domain specific binding | up | 60 | 7.00E-14 |
| positive regulation of signal transduction | up | 58 | 8.70E-14 |
| Genes involved in DNA Replication | up | 32 | 9.20E-14 |
| cell cycle | up | 67 | 3.00E-13 |
| Genes involved in Adaptive Immune System | up | 56 | 3.20E-13 |
| Focal adhesion | up | 31 | 5.60E-13 |
| kinase binding | up | 44 | 5.50E-12 |
| cytoskeleton organization | up | 54 | 5.60E-12 |
| neuron differentiation | up | 52 | 5.80E-12 |
| basement membrane | up | 19 | 7.50E-12 |
| regulation of cell migration | up | 36 | 1.10E-11 |
| MAPKinase Signaling Pathway | up | 20 | 1.40E-11 |

| hiL genes (human) |  |  |  |
| --- | --- | --- | --- |
| Biogroup name | direction | Common genes | p-value |
| chromosome | down | 65 | 1.30E-32 |
| response to DNA damage stimulus | down | 60 | 4.80E-32 |
| Genes involved in Cell Cycle | down | 50 | 3.60E-31 |
| Genes involved in Cell Cycle, Mitotic | down | 43 | 1.20E-30 |
| Genes involved in DNA Replication | down | 34 | 2.00E-30 |
| interphase | down | 47 | 8.20E-30 |
| interphase of mitotic cell cycle | down | 46 | 5.10E-29 |
| Genes involved in Mitotic M-M/G1 phases | down | 29 | 2.60E-26 |
| Cell cycle | down | 25 | 7.60E-23 |
| DNA repair | down | 40 | 2.30E-22 |
| wound healing | down | 50 | 7.40E-22 |
| response to ionizing radiation | down | 21 | 1.20E-20 |
| nuclear division | down | 35 | 8.00E-20 |
| mitosis | down | 35 | 8.00E-20 |
| cell division | down | 38 | 2.20E-19 |
| G1/S transition of mitotic cell cycle | down | 26 | 1.70E-18 |
| cardiovascular system development | down | 46 | 3.20E-18 |
| circulatory system development | down | 46 | 3.20E-18 |
| S phase | down | 22 | 3.80E-18 |
| blood coagulation | down | 41 | 6.10E-18 |

Among the mouse hiI genes, 4 out of the top 20 biogroups were associated with the nucleus (e.g., genes involved in the cell cycle and genes involved in DNA replication) (Table 1a). Among the human hiI genes, 15 out of the top 20 biogroups were associated with the nucleus (e.g., genes involved in the cell cycle, chromosomes, and response to DNA damage stimulus) (Table 1b). It is noteworthy that there is little overlap in the top 20 biogroups for the hiM and hiI genes (Table 1a, 1b). Thus, the biogroups related to the hiM and hiI genes are likely to be functionally different.

We also compared datasets from the DG of typically developing infants with datasets from rat DG at three different timepoints after seizure induction by injection of pilocarpine or kainite (day 1, day 3, and day 10), and performed principal component analysis on the changes in mouse hiM/hiI genes at different timepoints (Figure S2a, S2b; Supplementary Results). The time-course of changes in the mouse hiM genes after seizure induction was different from the time-course of changes in the mouse hiI genes. In addition, we conducted a spatial pattern analysis of the mouse hiM/hiI genes, which indicated that their protein products have slightly different patterns of subcellular localization (Figure S2c; Supplementary Results). The mouse hiM genes tend to be strongly expressed at the plasma membrane, with expression changes stabilizing by the third day after seizure induction. By contrast, the hiI genes tend to be expressed in the nucleus and changes in expression after seizure induction are slower to stabilize. Together, these results indicate that the hiM/hiI genes have different spatiotemporal patterns of changes in expression.

### Gene expression patterns in patients can be characterized in terms of hiM/hiI genes

Next, we investigated whether and to what extent the expression changes in maturation-related genes induced by hyperexcitation overlap with gene expression patterns in various neuropsychiatric disorders. As above, we evaluated similarities between the changes in gene expression patterns in different groups using overlap *P*-values calculated by Running Fisher algorithm (Figure 2a). Similarity indexes for each comparison were defined as the -log of the overlap *P*-values with hiM or hiI genes, denoted by hiM-index or hiI-index, respectively. High values in hiM-/hiI-index represent that there is large overlap between the dataset analyzed and hiM/hiI genes. We obtained the hiM-/hiI-indexes for the datasets from human patients and plotted them in two-dimensional (2-D) space to show the extent of overlap between datasets and hiM/hiI genes (Figure 2a).

**Figure 2.**
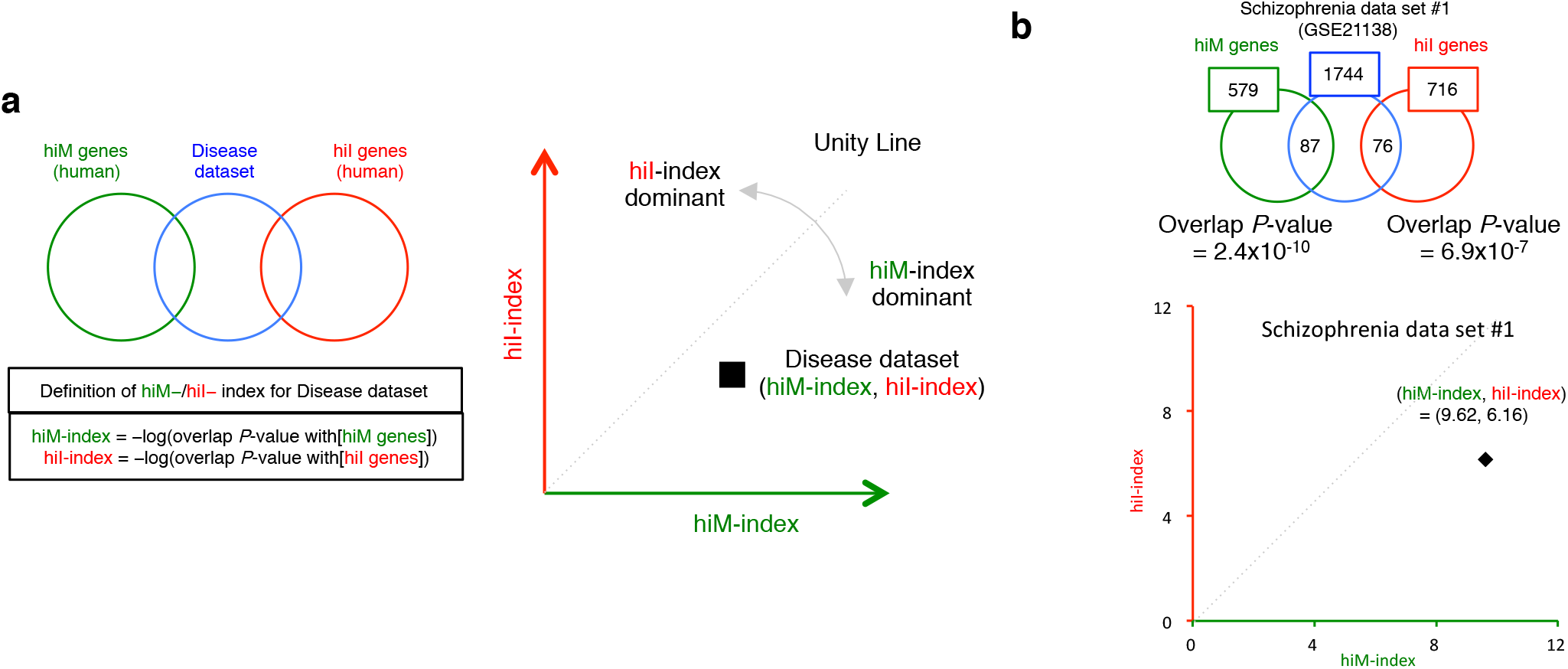
Overview of the two-dimensional analysis (2-D analysis) for disease datasets. (a) Genes with expression changes in the disease datasets are compared with the hiM and hiI gene groups. The hiM- and hiI-indexes were defined as the -log of the overlap *P*-values with the hiM and hiI genes. The gene expression patterns of the disease datasets are plotted in two-dimensional coordinates, in which the x-/y-axes are defined by the hiM-/hiI-indexes. Each dataset is characterized as hiM- or hiI-dominant by the ratio of hiM-/hiI-indexes, and the degree of the hiM-/hiI-dominance are evaluated by deviation from the unity line. Distance of each dataset from the origin show the degree of overlap with hiM-/hiI-genes. (b) 2-D analysis applied to a dataset of postmortem brains (prefrontal cortex) from patients with schizophrenia (schizophrenia dataset #1). The expression levels of 1744 genes were significantly changed in this disease dataset. Of these, 87 and 76 genes overlap with the hiM/hiI genes. The overlap *P*-values between the disease dataset and the hiM/hiI genes were 2.4 × 10^-10^ and 6.9 × 10^-7^. The hiM- and hiI-indexes for the disease dataset were therefore 9.62 and 6.16, indicating that this dataset is hiM-dominant. The results of the 2-D analysis for this dataset are plotted in the two-dimensional coordinates defined by the hiM- and hiI-indexes.

We initially performed this 2-D analysis on a dataset containing the expression profile of the prefrontal cortex in the postmortem brains of patients with schizophrenia (schizophrenia dataset #1: details in Table S2) (Figure 2b). The expression of 1744 genes differed between patients and healthy controls (significance level of 0.05). The numbers of hiM and hiI genes with altered expression in schizophrenia dataset #1 were 87 and 76, respectively, and the overlap *P*-values were 2.4 × 10^-10^ and 6.9 × 10^-7^. The hiM-index and hiI-index for this dataset were 9.62 (= -log(2.4×10^-10^)) and 6.16 (= -log(6.9×10^-7^)). This result corresponds to a point in 2-D space (Figure 2b). Points that fall below the unity line (dashed line) indicate datasets in which changes in hiM genes are dominant, whereas points above the unity line indicate datasets with dominant changes in hiI genes. The angle from unity line indicates the degree of the hiM-/hiI-dominance. The same analysis was performed for other schizophrenia datasets (schizophrenia datasets #2-#16), including ones obtained from different areas of the postmortem brain and from cultured neurons derived from the iPSCs of patients. Scatter plots of the results from the schizophrenia datasets are shown in Figure 3a. Thirteen out of sixteen points were below the unity line. Most of the schizophrenia datasets exhibited hiM-index dominant patterns, showing high hiM-index values and low hiI-index values (Figure 3a).

**Figure 3.**
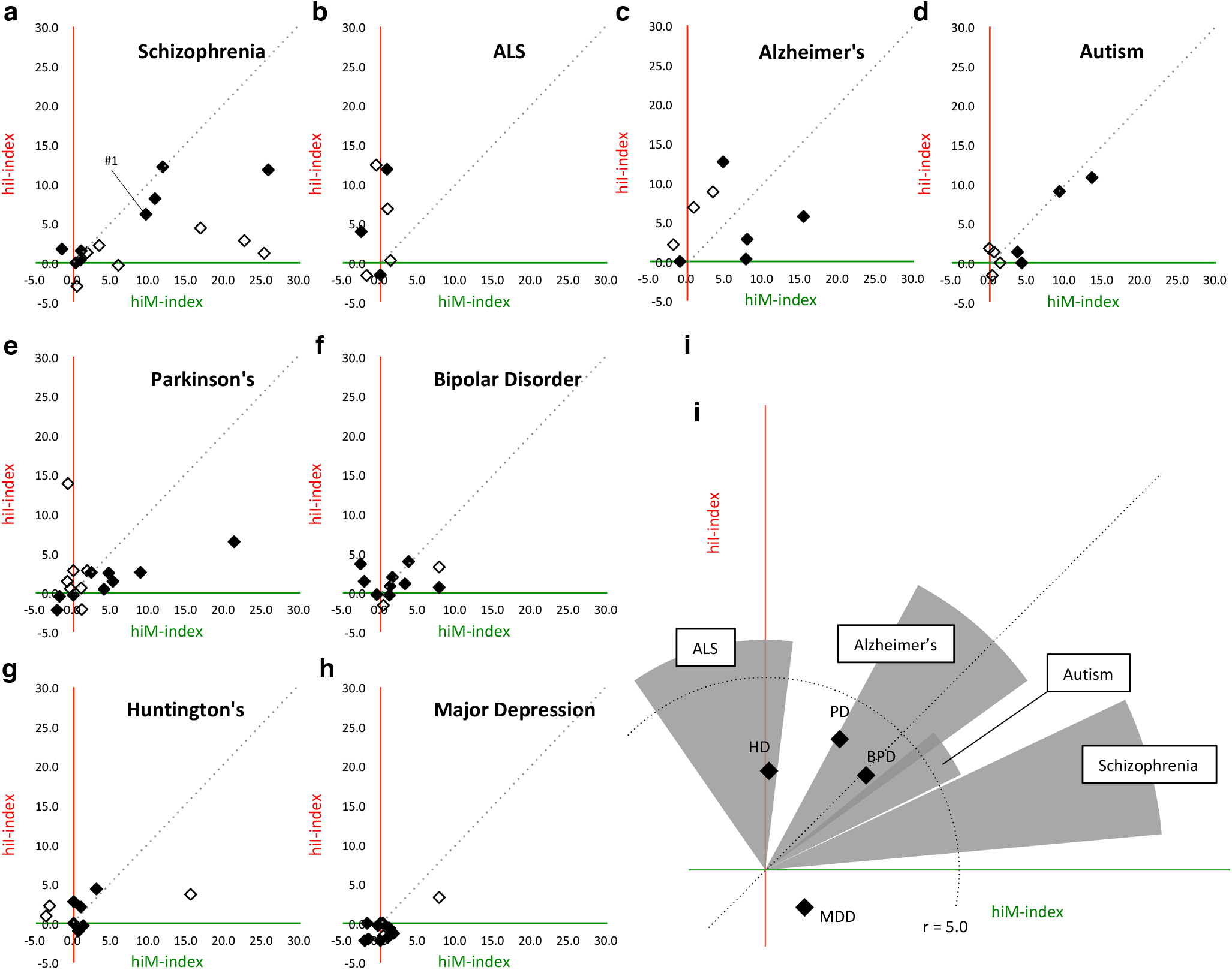
Two-dimensional analysis (2-D analysis) for disease datasets from various neuropsychiatric disorders. Each point corresponds to the result from one independent study. (a-h) Results of the 2-D analysis of datasets for schizophrenia (a), ALS (b), Alzheimer’s disease (c), autism (d), Parkinson’s disease (e), bipolar disorder (f), Huntington’s disease (g), and major depression (h). Filled points indicate datasets from the postmortem brain or spinal cord (ALS) of patients, and open points indicate those from cultured neural cells from patient iPSCs. (i) The distribution patterns of hiM- and hiI-index for all diseases analyzed. The extent of the changes in hiM-/hiI-indexes is assessed by the average distance of all datasets in each disease from origins. Four diseases whose average distance from origin are over 5.0 are shown as circular sectors, and others are shown as points. The radii of circular sector indicate the average distance of all datasets in each disease from origins, and the central angles of circular sector are average deviation ± sem from unity line. Each point indicates their average distance from origin and average deviation from unity line.

We extended the same analysis to 80 disease datasets from seven other neuropsychiatric diseases (amyotrophic lateral sclerosis (ALS), Alzheimer’s disease (AD), autism spectrum disorder (ASD), Parkinson’s disease (PD), bipolar disorder (BPD), Huntington’s disease (HD), and major depressive disorder (MDD); Table S2). Results of each dataset are shown in Figure 3b-3h. Overall distribution patterns of each disease are shown in Figure 3i. The ALS datasets tended to show higher hiI-index values than hiM-index values, indicating hiI-index-dominant pattern (Figure 3b). The AD datasets showed different patterns in the hiM-/hiI-index depending on the type of sample; datasets from the postmortem brains of patients with AD tended to show high values only in the hiM-index, and datasets from patient iPSCs tended to show high values only in the hiI-index (Figure 3c). Datasets from ASD did not show any dominant patterns either in the hiM-index or hiI-index (Figure 3d). Most datasets from patients with PD, BPD, HD, and MDD did not show pronounced values in the hiM-index or the hiI-index (Figure 3e, 3f, 3g, 3h).

Thus, the 2-D analysis revealed that some neuropsychiatric diseases have characteristic patterns in the hiM-/hiI-indexes: for example, most datasets from patients with schizophrenia exhibited a higher hiM-index than hiI-index, whereas ALS datasets showed hiI-index-dominant pattern (Figure 3i). Meanwhile, different diseases sometimes show similar changes in the hiM- or hiI-index; for example, some of the schizophrenia, ASD, and AD datasets shared a high hiM-index, and some of the ALS, and AD datasets shared a high hiI-index. The other four diseases—PD, BPD, HD, and MDD—did not show pronounced changes in hiM-/hiI-indexes, suggesting that these diseases may not share endophenotype as pseudoimmaturity inducible by neural hyperexcitation. These results raise the possibility that there are patterns of gene-expression perturbations that are shared and distinct across these neuropsychiatric disorders.

### Genetic and environmental risk factors induce changes in the pattern of expression of hiM/hiI genes

Previous studies suggest that many genetic risk and environmental factors, such as seizure, hypoxia, and infection, contribute to the development of neuropsychiatric disorders^2, 49, 50^. We next applied the 2-D analysis technique to datasets from genetic animal models of disorders and from animals that experienced risk events.

First, we obtained publicly available datasets for mice that had experienced putative risk events for schizophrenia, bipolar disorders, and Alzheimer’s disease, including seizure (#1: GSE49030, #2: GSE4236)^51, 52^, ischemia (#1: GSE32529, #2: GSE35338)^1–3^, and infection (mimicked by CpG; GSE32529)^53, 54^. All the studies used here included datasets for different time points after the risk event; hence, we were able to examine the time-course of changes in the hiM- and hiI-indexes to reveal the short- and long-term effect of risk event on expression patterns of hiM/hiI genes. The results showed that datasets from mouse hippocampus treated with kainite, seizure-inducing drug, exhibited time-course changes in the hiM- and hiI-indexes: the hiM-index tended to be dominant in the early stage after seizure induction, and the hiI-index became more dominant in the late stages (Figure 4a: seizure #1). The results from other datasets on seizure, ischemia, and infection showed roughly similar time-course pattern changes in the hiM- and hiI-indexes as those observed in seizure dataset #1, being relatively hiM-index-dominant in the early stage, and then relatively hiI-index-dominant in the later stage (Figure 4a: seizure, ischemia, and CpG infection). These results indicate that different types of putative risk events for neuropsychiatric disorders induce roughly similar time-course changes in the expression of maturation-related genes induced by neural hyperexcitation.

**Figure 4.**
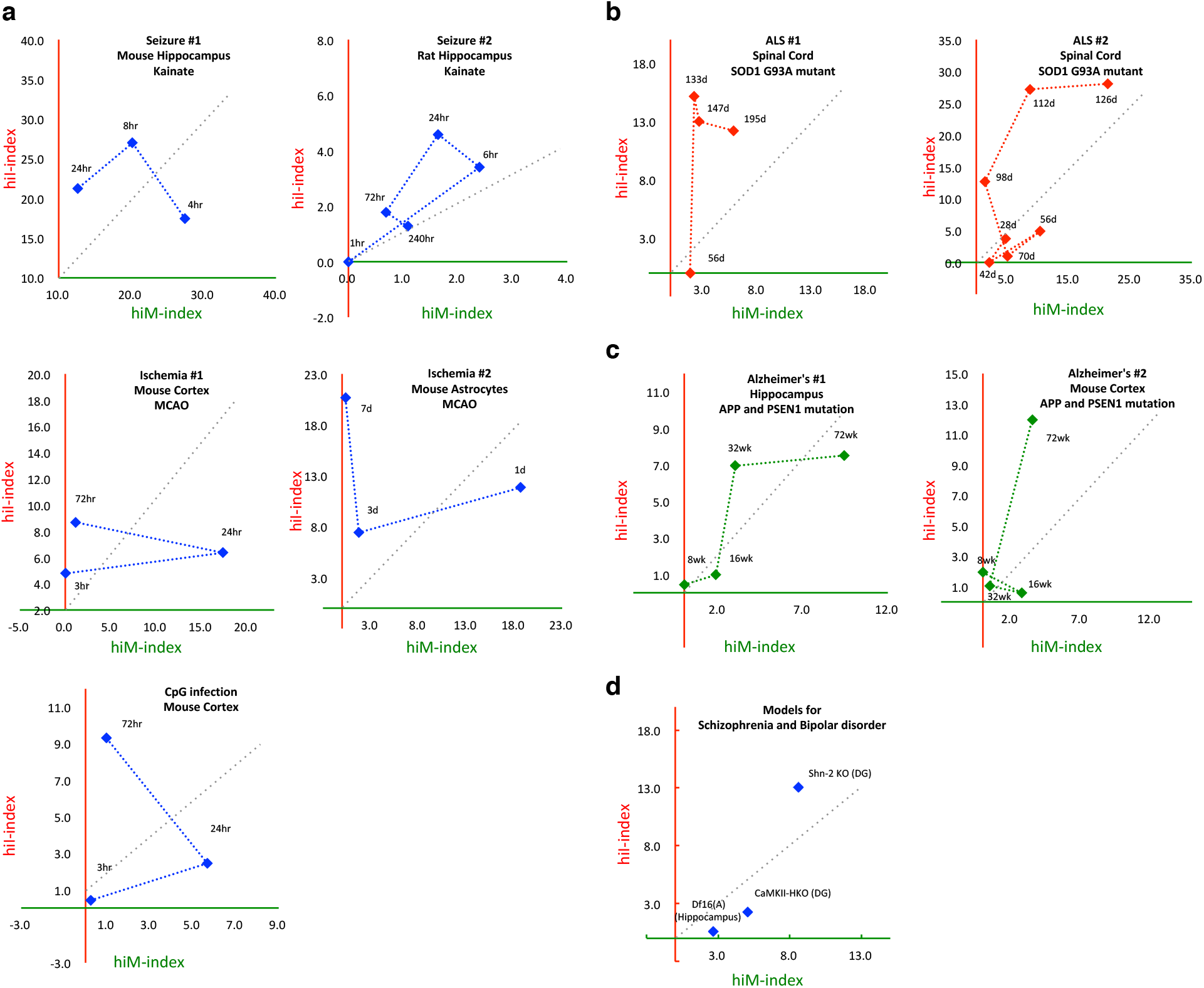
Time-dependent changes in the hiM-/hiI-indexes in animals subjected to various putative risk events for neuropsychiatric disorders and in genetic mouse models of schizophrenia, bipolar disorder, ALS, and Alzheimer’s disease. (a) Pattern of changes in the hiM- and hiI-indexes in mouse and rat hippocampus after treatment with kainite (seizure #1 (GSE1831) and seizure #2 (GSE4236)), in mouse cortex and astrocytes after middle cerebral artery occlusion (MCAO; ischemia #1 (GSE32529), ischemia #2 (GSE35338)), and in mouse cortex after CpG infection (GSE32529). (b) Pattern of changes in the hiM- and hiI-indexes of the spinal cord of an ALS mouse model with the SOD1(G93A) mutation. (c) Pattern of changes in the hiM- and hiI-indexes of the hippocampus and cortex of an Alzheimer’s disease mouse model with mutations in APP and PSEN1. (d) hiM- and hiI-indexes in mouse models of schizophrenia and bipolar disorder.

Next, we obtained datasets from animal models with a genetic risk of a neurodegenerative disease: mice with transgenic expression of a G93A mutant form of human SOD1, as a model of ALS (#1: GSE46298, #2: GSE18 5 97)^55, 56^; transgenic mice with mutant human amyloid precursor protein (APP) and presenilin1 (PSEN1) genes, which cause familial Alzheimer’s disease (#1: GSE64398, #2: GSE64398)^57^; and Df16(A) heterozygous mice carrying a chromosome 16 deletion syntenic to human 22q11.2 microdeletions, as a model of schizophrenia (GSE29767)^58^. We also obtained datasets from Schnurri-2 (Shn-2) knockout mice as a model of schizophrenia^12^ and intellectual disability^14, 59, 60^ and from mice with heterozygous knockout of the alpha-isoform of calcium/calmodulin-dependent protein kinase II (alpha-CaMKII+/-)^10, 13^ as a model of bipolar disorder. We performed 2-D analysis on these datasets and evaluated the changes in the hiM-/hiI-indexes of these model mice. Both datasets from transgenic mice with a SOD1 (G93A) mutation exhibited a higher hiI-index than hiM-index in the later stages of disease progression (Figure 4b). These hiI-index-dominant patterns were also observed in the results derived from human patients with ALS (Figure 3b). In the mice with mutant human APP and PSEN1, both the hiM- and hiI-indexes increased in dataset from hippocampus and only hiI-index increased in dataset from cortex during the course of disease progression (Figure 4c). These patterns are neutral or hiI-index-dominant. These patterns partially mimic the results from human patients with Alzheimer’s disease (Figure 3c). The Df16(A) heterozygous mice and alpha-CaMKII+/- mice showed hiM-index-dominant patterns, which are similar to results from human patients with schizophrenia (Figure 4d). Shn-2 KO mice showed high values for both the hiM- and hiI-indexes (Figure 4d). Thus, the results from the 2-D analysis of animal models are to some extent consistent with the results from human patients, indicating that these model mice share similar patterns of pseudoimmaturity induced by neural hyperexcitation with those of human patients.

## Discussion

In this study, we demonstrated that neural hyperexcitation induces changes in the pattern of gene expression in the DG that are significantly similar to the immature hippocampus of typically developing human fetuses. From the pool of genes, we identified two groups of genes, and found that these are shared by multiple neuropsychiatric disorders, such as schizophrenia, Alzheimer disorders, and ALS.

Many of the datasets from patients with schizophrenia and from the postmortem brains of patients with Alzheimer’s disease exhibited hiM-index-dominant pattern changes. The hiM genes include a GABA receptor, voltage-dependent calcium channel, glutamate receptor, and voltage-dependent sodium channel (Table S1). These genes have been reported to be implicated in the pathological changes in the brains of patients with schizophrenia and Alzheimer’s disease^61–64^. Thus, many of the synaptic genes that changed in the brains of patients with schizophrenia or Alzheimer’s disease could be genes whose expression increases during maturation and decreases with neural hyperexcitation. Although reductions in the expression of some synaptic genes in these disorders are well documented, our results are the first to raise the possibility that neuronal hyperexcitation may also induce reductions in such synaptic molecules.

Most of the datasets from patients with ALS and Alzheimer’s disease exhibited hiI-index-dominant patterns. The hiI genes include DNA methyltransferase, cyclin, cyclin-dependent kinase, integrin beta 3 binding protein, and tumor protein p53 (Table S1). These genes are known to be important in chromosomal modification and DNA repair, and abnormal functions of these systems have been observed in patients with ALS and Alzheimer’s disease^65–69^. Thus, some of the genes that are considered to be important in the development of these disorders are immaturity-related genes, whose expressions decrease during maturation and can be increased by neural hyperexcitation.

As for the datasets from patients with PD, BPD, HD, and MDD, most of them did not show significant overlap either hiM or hiI genes, indicating that there might not be pathological changes of pseudoimmaturity inducible by neural hyperexcitation in the datasets of these four diseases. Thus, we suggest that gene expression analysis based on the findings derived from shared endophenotypes are helpful to conduct transdiagnostic characterization of neuropsychiatric disorders.

Our study has some limitations. First, the number of available datasets was limited. All the datasets except the one for mouse development were obtained from the BaseSpace Correlation Engine. On this platform, vast numbers (over 21,000) of complex biological and clinical datasets are available. Even though we used all the gene expression dataset hits from our keyword query to avoid sampling bias, the number of datasets was still small, from 8 datasets for ALS to 16 datasets for schizophrenia. Further accumulation of the studies will improve the reliability of our results. Another limitation is that the datasets used in this study are from different types of sample, including various central nervous system areas, such as the hippocampus, prefrontal cortex, striatum, and the spinal cord. The gene expression abnormalities in patients could differ depending on the brain area^48^. We also used datasets from cultured neurons differentiated from the iPSCs of human subjects, and it is controversial whether the pattern of gene expression in these neurons is comparable with that of neurons in the patients’ brains^70, 71^. It is also possible that the altered gene expression in the postmortem brains is due to the effects of medication rather than pathological changes from the disease itself^72^. Other conditions that were not controlled in this study include the age at death, storage conditions of the samples, genetic background of animals, and animal housing conditions. For these reasons, we need to be careful in interpreting the results of the analyses. It is noteworthy, however, that despite the variety of sample types used, we were able to identify some shared and distinct patterns of gene expression.

Recent attempts such as RDoC initiative have tried to reclassify psychiatric disorders according to biological domains (e.g. genes, neural circuits, behavior)^7^. While Gandal *et al* conducted non-biased coexpression analyses^8^, in this study, we utilized gene groups which were derived from the findings based on the studies of animal models of neuropsychiatric disorders. Characterization by these gene groups enabled us to extract novel biological features of some neuropsychiatric disorders which are related to pseudoimmaturity inducible by neural hyperexcitation. Detecting such domains that extract the biological features of each neuropsychiatric disorder will move this diagnostic framework forward, from criteria based on signs and symptoms to those including biological dimensions.

In conclusion, biological domain, which is pseudoimmaturity inducible by neural hyperexcitation, are common endophenotype among several neuropsychiatric disorders. Future studies are needed to find translational indices that correspond to these features and can be applicable to human patients for better diagnosis of these neuropsychiatric disorders. Our findings here may promote the development of novel biomarkers, leading to better diagnosis of neuropsychiatric disorders.

## Methods

### Microarray experiments to examine mouse DG development

Wild-type mouse DGs were sampled at postnatal day 8, 11, 14, 17, 21, 25, 29 (C57BL/6J × BALB/cA background; n = 5)^73^, and microarray experiments were performed as previously described^10^. The microarray data, including those used in this study, were deposited in the GEO database under accession number GSE113727. We also obtained a dataset for 33-week-old wild-type mice, which we previously reported (C57BL/6J × BALB/cA background) (GSE42778)^12^. We integrated these two datasets into one to construct the dataset for the development of wild-type mouse DG used in this study (P8 versus adult, fold change > 1.2, *P* < 0.05).

### Data collection and processing

Except for the mouse DG developmental dataset mentioned above, the 91 gene expression datasets used in this study were obtained from publicly available databases (listed in Table S2). All gene expression datasets were analyzed with the BaseSpace Correlation Engine (BSCE; formally known as NextBio) (https://japan.ussc.informatics.illumina.com/c/nextbio.nb; Illumina, Cupertino, CA), a database of biomedical experiments. BaseSpace is a repository of analyzed gene expression datasets that allows researchers to search expression profiles and other results^44^. The datasets registered in BaseSpace undergo several preprocessing, quality control, and organization stages. Quality control ensures the integrity of the samples and datasets and includes evaluations of pre- and postnormalization boxplots, missing value counts, and *P*-value histograms (after statistical testing) with false-discovery rate analysis to establish whether the number of significantly altered genes is larger than that expected by chance. Other microarray data processing was performed in MAS5 (Affymetrix, Santa Clara, CA, USA)^44^.

Genes with a *P*-value < 0.05 (without correction for multiple testing) and an absolute fold change > 1.2 were included in the differentially expressed gene datasets. This sensitivity threshold is typically the lowest used with commercial microarray platforms and the default criterion in BaseSpace analyses^44^. All data from the Affymetrix GeneChip series were downloaded from the NCBI GEO database. Affymetrix Expression Console software (specifically the robust multiarray average algorithm) was used to preprocess the data.

We used the expression values (on a log base-2 scale) to calculate the fold changes and *P*-values between two conditions (infants–adults and patients–healthy controls). To determine the fold changes, the expression values of the probes/genes in the test data sets were divided by those of the control data sets. If the fold change was < 1.0, these values were converted into the negative reciprocal or −1/(fold change). Genes with an absolute fold change > 1.2 and a *t*-test *P*-value < 0.05 were imported into BSCE according to the instructions provided by the manufacturer. The rank order of these genes was determined by their absolute fold change. We compared the signatures in two given gene sets using BSCE. All statistical analyses were performed in BaseSpace and similarities between any two datasets were evaluated as overlap *P*-values using the Running Fisher algorithm^44^.

### Funding and Disclosures

Competing financial interests: K.T. and M.M. are employees of Astellas Pharma Inc. Other authors report no biomedical financial interests.

## Acknowledgments

We thank Wakako Hasegawa, Yumiko Mobayashi, Misako Murai, Tamaki Murakami, Miwa Takeuchi, Yoko Kagami, Harumi Mitsuya, Aki Miyakawa and other members of Miyakawa lab for their support. This work was supported by JSPS Grant-in-Aid for Scientific Research on Innovative Areas Grant Number 25116526, 15H01297 (“Microendophenotype”), 16H06462 (“Scrap & Build”), JSPS KAKENHI Grant Number 25242078, and grant from Astellas Pharma Inc.

## Author Contribution Statement

T.M. generated the data, performed the analysis, prepared all figures and wrote the manuscript. All authors reviewed the manuscript.

